# Complete genomes of DNA viruses in faecal samples from small terrestrial mammals in Spain

**DOI:** 10.1101/2024.11.12.623162

**Authors:** Jaime Buigues, Adrià Viñals, Raquel Martínez-Recio, Juan S. Monrós, Rafael Sanjuán, José M. Cuevas

**Affiliations:** Institute for Integrative Systems Biology (I2SysBio), Universitat de València and Consejo Superior de Investigaciones Científicas, València, Spain; Institut Cavanilles de Biodiversitat i Biologia Evolutiva, Universitat de València, València, Spain; Department of Genetics, Universitat de València, València, Spain

**Author notes:** Competing interests: The authors declare no competing interests.

**Keywords:** DNA viruses, metagenomics, viral emergence, viromics, zoonotic viruses

## Abstract

Viromics studies are allowing us to understand not only the enormous diversity of the virosphere, but also the potential threat posed by emerging viruses. Regarding the latter, the main concern lies in monitoring the presence of RNA viruses, but the zoonotic potential of some DNA viruses, on which we have focused in the present study, should also be highlighted. For this purpose, we analysed 160 faecal samples from 14 species belonging to three orders of small terrestrial mammals (i.e. *Rodentia, Lagomorpha* and *Eulypotyphla*). This allowed us to identify a total of 25 complete or near-complete genomes belonging to the families *Papillomaviridae, Polyomaviridae, Adenoviridae, Circoviridae* and *Genomoviridae*, 18 of which could be considered new species or types. Our results provide a significant increase in the number of complete genomes of DNA viruses of European origin with zoonotic potential in databases, which are at present clearly under-represented compared to RNA viruses.

## 1. Introduction

Metagenomics has accelerated the rate of virus discovery in clinical and environmental samples [1,2], thus increasing our understanding of the global virosphere [1]. In addition, metagenomics can be used as a promising tool for the prevention of zoonotic diseases through early detection of viral reservoirs, thereby reducing the risk of zoonotic outbreaks [3,4]. For this purpose, a commonly used non-invasive method is the analysis of faecal samples from wild animals, which has led to the identification of numerous viruses with zoonotic potential [5–7].

In the search for viruses with zoonotic potential, bats are considered the most important mammalian reservoir [8]. Several zoonotic outbreaks caused by coronaviruses [9,10] and paramyxoviruses [11,12], among others, have been associated with bats. Other mammals, however, can also act as potential viral reservoirs, with rodents being the best candidate, as they are the most specious mammalian group [13]. For example, dozens of zoonotic pathogens have been identified in rodents [14] and cause diseases in humans, such as hantavirus pulmonary syndrome [15] and Lassa fever [16].

Several rodent species live in close contact with humans, increasing the zoonosis risk. For instance, the house mouse has promoted global transmission of viruses to sympatric species [17]. Despite their clear role as viral reservoirs, studies in rodents have not attracted the same interest as those based on bats. This is evident when comparing the number of viral sequences deposited in bat-related databases such as DBatVir (http://www.mgc.ac.cn/DBatVir/), which had roughly 21,000 sequences in September 2024, with those deposited in the rodent database (DRodVir: http://www.mgc.ac.cn/DRodVir/), which included over 14,000 sequences. It is worth noting that most of the sequences deposited in both databases are partial. In addition, analogous to DBatVir, there is also a strong bias towards the description of RNA viruses in rodents, which account for 90% of the sequences deposited in DRodVir. This is probably because RNA viruses have an increased zoonotic potential overall [18], although many DNA viruses are also of particular concern [19].

Despite a quarter of the sequences deposited in DRodVir are from Europe, only 34 sequences correspond to Spanish origin, 5 of which are DNA viruses. In order to provide more information on potential reservoirs of DNA viruses in Spain, we have carried out a metagenomic analysis to characterise the DNA fraction of 160 faecal samples from fourteen species belonging to three different orders (i.e. *Rodentia, Lagomorpha* and *Eulypotyphla*). Overall, the assembly of viral reads obtained has allowed the recovery of 25 complete or nearly complete metagenome-assembled viral genomes (MAVGs) belonging to groups with zoonotic potential, 18 of which represent novel DNA virus species or types.

## 2. Results and discussion

### 2.1. Preliminary analysis and global overview

Fecal saliva samples were collected from 160 individuals of 14 different species (**Table S1**). Preliminary studies of the samples collected suggested the presence of potential inhibitors at some stage leading up to the massive sequencing data (Results not shown). Some samples could contain substances promoting the degradation of nucleic acids/viral particles or inhibiting extraction/sequencing stages, such as urine [20] or dietary components [21]. To test this, vesicular stomatitis virus (VSV) was spiked into an aliquot of each sample before starting processing (see Materials and Methods). Subsequently, the recovery of VSV was quantified in plaque assays and samples showing >90% reduction of VSV viability compared to positive controls without faecal content were discarded. This potential inhibitory effect shown by plaque assays was further confirmed in some samples by quantitative PCR (qPCR) of VSV (Results not shown). As a result, only 76 samples of 9 different species from 6 Spanish provinces were analyzed (**Figure 1** and **Table S1**). At this point, it should be noted that the presence of potential inhibitors was not previously observed in bat viromics studies following exactly the same sample processing [22,23], although these were fresh samples, most likely free of urine residues, in addition to the inherent differences in diet composition. In any case, this underlines the convenience of evaluating the presence of potential inhibitors as a preliminary step in viromics studies.

**Figure 1.**
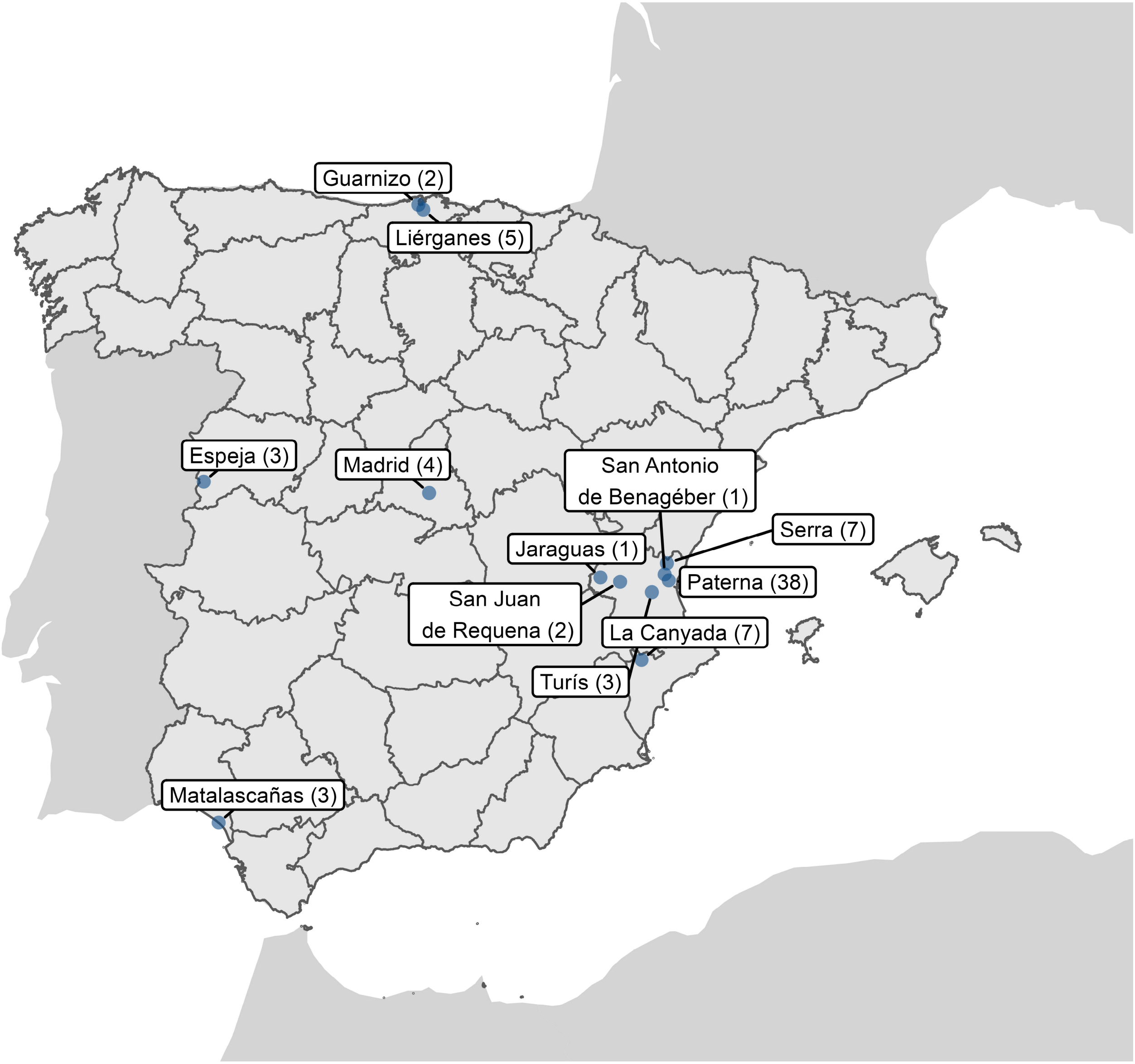
Sampling sites of the sequenced individuals. The number of individuals captured in each location is indicated in parentheses. This map was created using R software (https://www.R-project.org/).

Selected samples were processed in 13 pools, each containing exclusively samples belonging to the same species (**Table S1**). Illumina sequencing of the DNA samples generated between 15 and 32 million raw reads per pool (**Table S2**). After the first filtering step, roughly 30% of the reads were removed, most of which were probably PCR duplicates [23]. Filtered reads were de novo assembled, resulting in 72,485 viral contigs larger than 1 kb, of which 1,804 were complete or nearly complete genomes according to CheckV [24], although a vast majority were taxonomically classified as bacteriophage (**Table S3**). However, from the set of metagenome-assembled viral genomes (MAVGs), we focused on those associated with vertebrate viral families (i.e. *Papillomaviridae, Polyomaviridae, Adenoviridae* and *Circoviridae*), as well as the poorly studied *Genomoviridae* family, which has been associated to multiple virome studies [23,25,26]. In total, 25 MAVGs belonging to these families were found in 4 pools of two rodent and two shrew species (**Figure 2** and **Tables S1** and **S4**). One single MAVG corresponded to a linear genome virus (i.e. adenovirus), while 21 of the other 24 MAVGs (i.e. 87,5%) showed terminal redundancy and could therefore be considered as complete genomes. In a recent bat faecal viromics study, exactly the same experimental procedure was followed and twice as many MAVGs related to DNA viruses with the potential to infect vertebrates were identified [23]. However, it should be noted that the number of bat samples and species analysed was also approximately twice as high as in the present work. Therefore, the results obtained in both papers are consistent, regardless of the differences in the origin of the samples. This suggests that the methodology employed is appropriate, in general terms, except for the precautions to be taken in terrestrial mammal samples due to the presence of potential inhibitors, as mentioned above. It seems not strictly necessary, thus, to implement more complex procedures involving ultracentrifugation [27] or probe capture assays [28].

**Figure 2.**
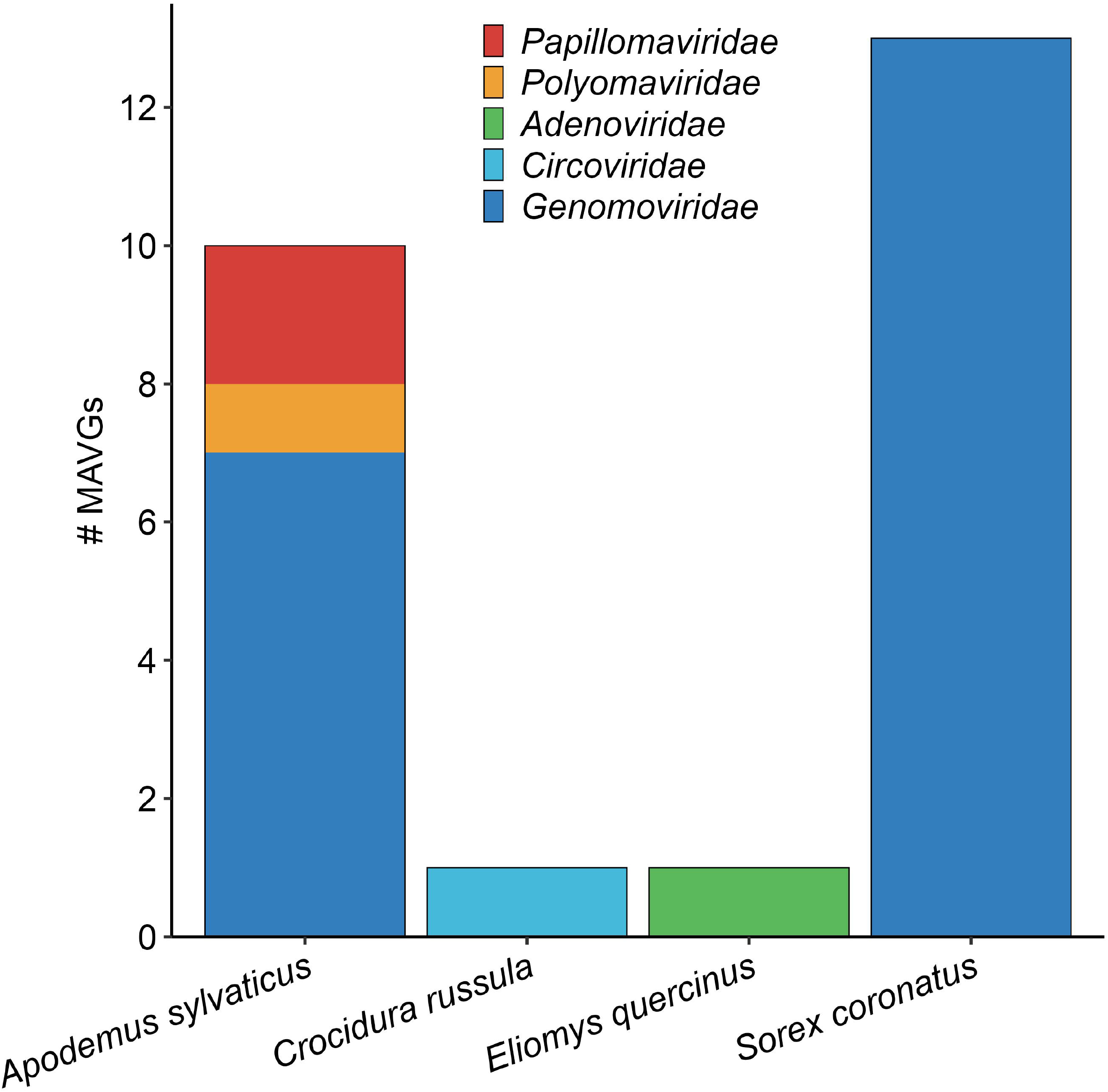
Distribution of MAVGs per species/pool. Viral families are shown in different colours.

To estimate the reliability of the MAVGs, each one was remapped with reads from its sequencing library, which showed a large variability ranging from a minimum of 121 reads (mean coverage depth 7.81) for MAVG12 to a maximum of 69,853 reads (mean coverage depth 252.62) for MAVG4 (**Table S4**). Based on BLASTn’s analyses, five of the MAVGs showed >85% sequence identity with previously described viruses at >90% coverage, while 20 corresponded to potential new viruses. In addition, four MAVGs (i.e. MAVG2, MAVG4, MAVG22, and MAVG25) had low coverages (i.e. <20%) with respect to the best Blast hit, and MAVG21 showed no hit. This could reflect the absence of sequences with sufficient genome-wide identity in databases. In these five cases, a BLASTp analysis using the largest inferred ORF showed a query coverage ranging from 88% to 100% and corroborated the previously assigned taxonomic classification. (**Table S4**). In the following, for the sake of simplicity, the description of the different MAVGs will be done separately for each virus family and those cases where new species can be considered according to the criteria established for each family will be detailed.

### 2.2. Novel papillomaviruses

MAVG1 and MAVG2 were classified as papillomaviruses. Both MAVGs were detected in fecal samples from *Apodemus sylvaticus* collected from a locality in Valencia (pool P38) (**Figure 2** and **Tables S1** and **S4**). Their genome sizes corresponded to those expected for a papillomavirus genome [29] and exhibited their typical organization, with early and late genes encoded on the same strand (**Table S5**) [30]. Detailed genome analysis of the two MAVGs identified several common amino acid motifs previously described in rodent papillomaviruses [31]. Both MAVGs contained two zinc-binding domains CX_2_CX_29_CX_2_C separated by 36 amino acids in the E6 protein. The ATP-binding domain (GX_4_GK(S/T)) was also present in the E1 protein of both papillomaviruses, whereas the leucine zipper domain (LX_6_LX_6_LX_6_L) was absent in the E2 protein of both MAVGs. In addition, the retinoblastoma protein (pRB) binding motif (LXCXE) was found only in MAVG2, as previously reported for other papillomavirus types associated with MAVG2 [31]. This pRB binding motif has been suggested to play an important role in the carcinogenesis process caused by papillomaviruses [32,33], although its presence is not strictly required for such an effect [34]. Finally, different regulatory elements were also detected in both MAVGs, including polyadenylation signals (AATAAA) and E1/E2 binding sites (E1BS and E2BS) [31,35,36]. However, the TATA box was not detected in MAVG2, probably because the contig lacked direct terminal repeats (DTRs) and thus is not a complete genome.

Papillomavirus taxonomy is based on nucleotide sequence identity across the L1 gene [37], thus sequences sharing >70% identity are considered viral variants of the same species. The sequences of these MAVGs were submitted to the International Animal Papillomavirus Reference Center (IAPRC), which validated the taxonomy and proposed a standard nomenclature. According to PaVE L1 Taxonomy Tool (https://pave.niaid.nih.gov), MAVG1 shared 76.95% sequence identity with Apodemus sylvaticus papillomavirus 1 (AsylPV1). In addition, a BLASTn analysis showed 97.9% genome-wide identity with a 100% coverage with Papillomavirus apodemus 7726 (Acc. BK066393.1), which was recently described [38]. Since this virus has not yet been included in the PaVE database, and following consultation with the IAPRC, MAVG1 was considered a new variant of AsylPV1 and no standard nomenclature was given. MAVG2 shared 71.86% L1 sequence identity with Rattus norvegicus papillomavirus 2 (RnPV2), in addition to 75.1% genome-wide identity with a query coverage of only 14%. Again, based on advice from the IAPRC, MAVG2 was considered a new type of the species to which RnPV2 belongs, and named as Apodemus sylvaticus papillomavirus 2 (AsylPV2). Complementing this taxonomic designation, a phylogenetic analysis using representative papillomaviruses assigned MAVG1 and MAVG2 to the genera *Pipapillomavirus* and *Iotapapillomavirus*, respectively (**Figure 3**). In both genera, the reference sequences present in the PaVE database belong to viruses from different rodent species and, in addition, the closest taxon to MAVG1 was isolated from the same species. In this sense, papillomavirus host-switching events are rare, but in some cases have been reported for genetically close related hosts [39].

**Figure 3.**
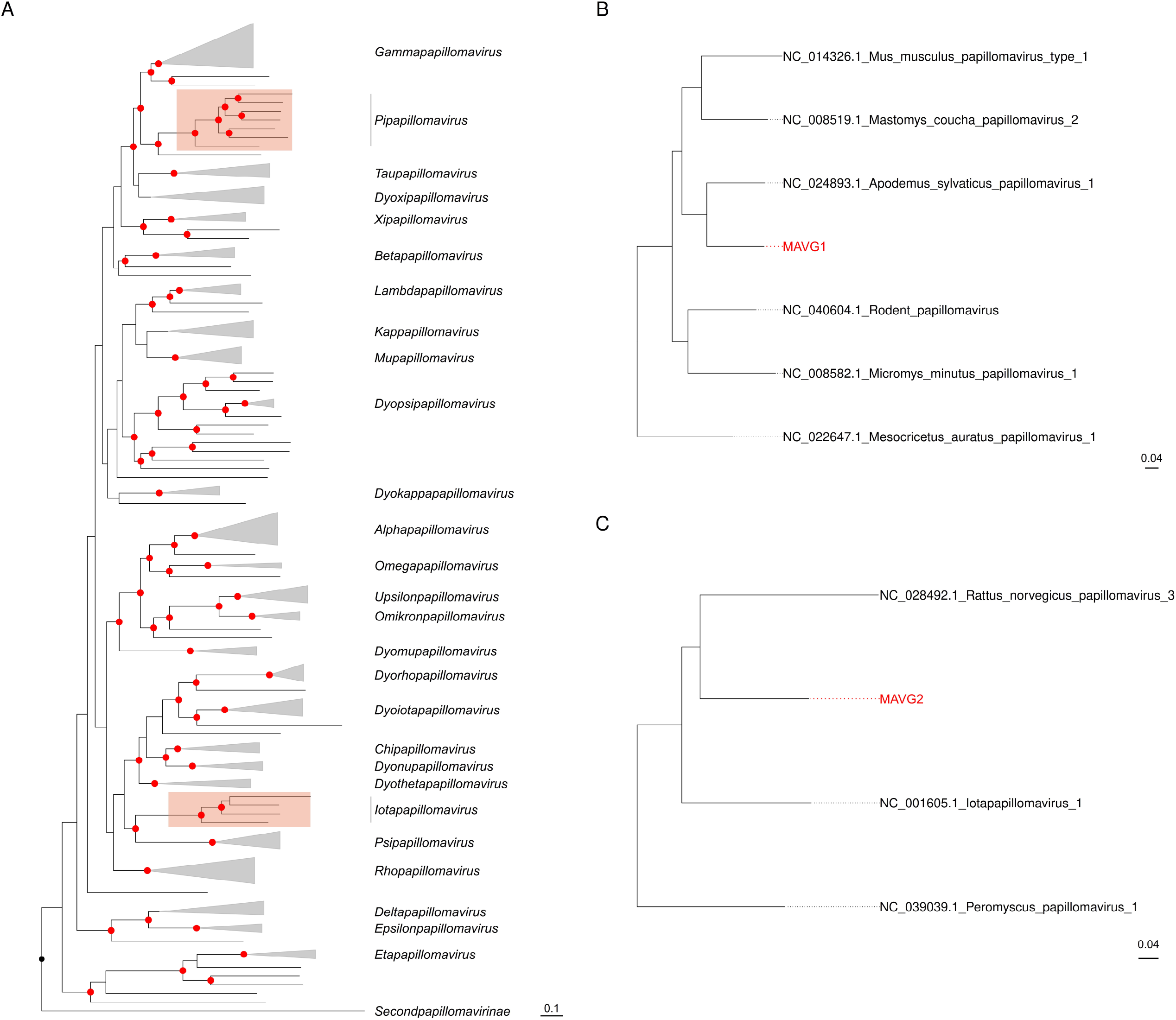
Maximum likelihood (ML) tree based on the concatenation of the E2-E1-L2-L1 gene nucleotide sequences from 206 representative papillomaviruses [23]. A) Full ML tree where clades corresponding to different genera are collapsed for clarity. B) Clade of the genus *Pipapillomavirus* containing MAVG1. C) Clade of the genus *Iotapapillomavirus* including MAVG2. Taxa are denoted by GenBank accession number and virus name, and novel MAVGs are marked in red. Phylogenetic analysis was done using substitution model GTR+F+I+G4. SH-aLRT and ultrafast bootstrap values higher than 80 and 95, respectively, are indicated in red circles. The tree is rooted at midpoint. The scale bar indicates the evolutionary distance in nucleotide substitutions per site.

### 2.3. A novel polyomavirus

MAVG3 was classified as polyomavirus and identified at the same pool as MAVG1 and MAVG2 (**Figure 2** and **Tables S1** and **S4**). The length of this MAVG (5.3 kb) corresponded to the expected size for a polyomavirus [40]. The genome organization of polyomaviruses consists of an early region encoding regulatory proteins, a late region for capsid proteins, and a non-coding regulatory region in between [40]. MAVG3 showed a genome organization typical of polyomaviruses, presenting four regulatory proteins (the large T antigen [LTAg], the small T antigen [STAg], the middle T antigen [MTAg] and the ALTO protein) and three capsid proteins (VP1, VP2 and VP3) (**Table S6**).

According to ICTV guidelines, two polyomaviruses belong to the same species if the amino acid sequence identity of the LTAg is higher than 70%. MAVG3 shared 95% sequence identity with *Alphapolyomavirus aflavicolis* (Acc. MG654477.1), so they could be considered variants of the same species. Despite this genetic proximity, the creation of a new species in polyomaviruses also depends on biological factors, such as host specificity, disease association or tissue tropism [41]. Consequently, considering that these two sequences have been isolated from different species of the genus *Apodemus*, MAVG3 could be considered a new species of polyomavirus. This taxonomic proposal was paralleled by a phylogenetic tree of the LTAg, which assigned MAVG3 to the genus *Alphapolyomavirus*, whose members infect primates, bats, rodents and other mammals (**Figure 4**) [41].

**Figure 4.**
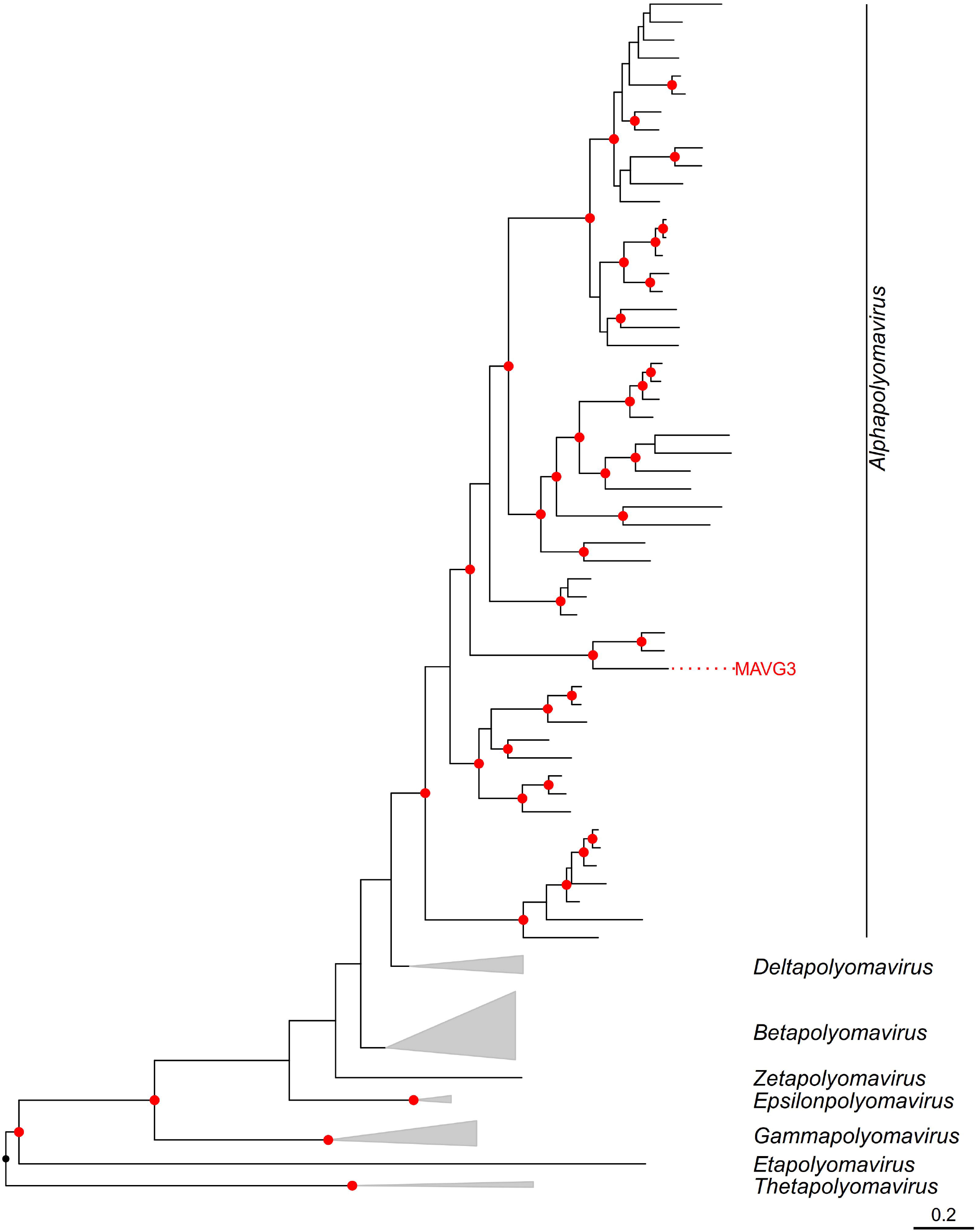
ML tree of the family *Polyomaviridae* using 135 RefSeq LTAg amino acid sequences [23]. Nodes are collapsed by genus and the novel MAVG is marked in red. Amino acid sequences were aligned using Clustal Omega v1.2.3 [48] and trimmed with trimAl v1.2rev59 [49] applying the *gappyout* parameter. Phylogenetic analysis was done using substitution model LG+F+I+G4. SH-aLRT and ultrafast bootstrap values higher than 80 and 95 respectively, are indicated in red circles. The tree is rooted at midpoint. The scale bar indicates the evolutionary distance in amino acid substitutions per site.

Following the demonstration of the oncogenic potential of a human polyomavirus, Merkel cell PyV (MCPyV), this family has attracted considerable interest [42]. ALTO protein is the only one that defines a monophyletic group of polyomaviruses [43], including MAVG3, and seems to play an important role in cancer development. Recent findings suggest that ALTOs evolved to suppress viral replication and promote viral latency and that MCPyV ALTO must be silenced to promote Merkel cell carcinoma development [44]. Therefore, the characterization of this protein in other species and its study in animal models may help to understand the mechanisms involved in oncogenesis. Globally, from an ecological perspective, the continuing discovery of polyomaviruses highlights the enormous diversity of this family, even on a small scale, as in the case of chimpanzee studies, where new polyomaviruses continue to be identified [45–47].

### 2.4. A novel adenovirus

An adenovirus genome, MAVG4, was assembled de novo in a pool of faecal samples from the rodent *Elyomis quercinus* (pool P41) (**Figure 2** and **Tables S1** and **S4**). MAVG4 showed the typical genome organization of *Mastadenovirus* genus, with a GC content of 51.4%, which is in the range described for mastadenoviruses [50]. This MAVG presented a putative E3 region characteristic of non-primate mastadenoviruses [51], which was 903 nucleotides long and lacked an ORF. Interestingly, the last 7.1 kb encoded ORFs including immunoglobulin protein domains (IPR036179 and IPR013783) and an OX-2 membrane glycoprotein-like domain (IPR047164), which is related to herpes viruses and has been associated with inflammation, autoimmunity control, hypersensitivity and spontaneous fetal loss [52]. In addition, MAVG4 presented inverted terminal repeats (ITR) of 28 bp at both ends of the genome and a total of 23 ORFs.

Two maximum likelihood (ML) trees were constructed using the DNA polymerase and hexon amino acid sequences, which are commonly used for taxonomic classification of mastadenoviruses [53]. These trees confirm that MAVG4 clustered with non-primate mastadenoviruses (**Figure 5**). However, species definition was more complex, as it depends on several factors, such as phylogenetic distance of DNA polymerase amino acid sequence, genome organization, or host range, among others [50]. At a genome-wide level, a BLASTn search showed 68.9% sequence identity and 4.0% query coverage with Taphozous bat adenovirus (Acc. PP711819.1) (**Table S4**). In addition, a BLASTp analysis of MAVG4 DNA polymerase showed a peak sequence identity of 65.4% and a 96% query coverage with Rhinolophus ferrumequinum adenovirus (Acc. WZK92861.1), reported in a Spanish bat. Also, a 75.9% peak sequence identity and 100% query coverage was observed with the hexon protein sequence of Lemur mastadenovirus (Acc. WGN96567.1). Consequently, considering that the most closely related sequences are highly distant and belong to non-rodent hosts, MAVG4 could be considered a new species associated with *Eliomys quercinus*. This small rodent is widely distributed in Spain, but its population is declining in Eastern Europe. For this reason, it is listed as Near Threatened on the IUCN Red List of Threatened Species (https://www.iucnredlist.org/species/7618/12835766), and the characterization of new viral pathogens associated with this species may contribute to its conservation.

**Figure 5.**
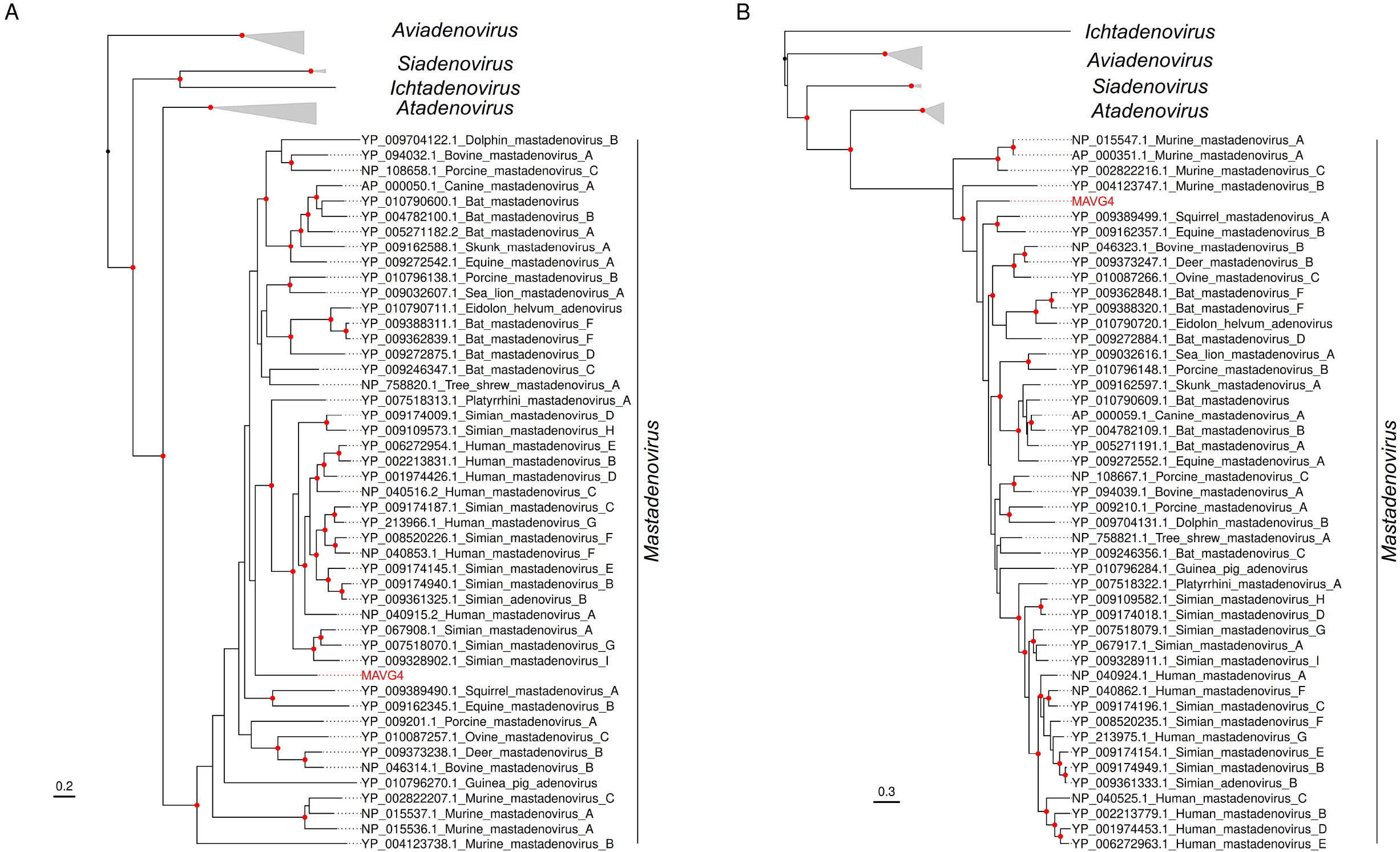
ML trees of the *Adenoviridae* family using DNA polymerase (A) and hexon (B) amino acid sequences from 73 representative members [23]. Taxonomic groups are collapsed by genus. Taxa are denoted by GenBank protein accession number and virus name, and the novel MAVG is labelled in red. Phylogenetic analyses were done using substitution models Q.pfam+F+I+G4 and Q.yeast+F+I+G4 for A and B trees, respectively. SH-aLRT and ultrafast bootstrap values higher than 80 and 95, respectively, are indicated in red circles. The tree is rooted at midpoint and the scale bar indicates the evolutionary distance in amino acid substitutions per site.

Adenoviruses have a relatively large genome size and viromics studies usually provide partial sequences [53–55]. However, the characterization of complete genomes, such as the one described in this work, is particularly relevant, given the prominent role of recombination in the genesis and emergence of adenoviral pathogens [56]. For example, intra- and interspecific recombination events have been described [57,58], in the latter case associated with adaptation to a new host. Related with this, although adenoviruses are usually species-specific, there is evidence of their zoonotic potential [56], not only for crossing the species barrier between humans and non-human primates [59], but also between humans and other animal species [60]. Regarding cross-transmission between different animal species, a notable example is given by the skunk adenovirus-1, which has been documented in seven mammalian families [61], strongly suggesting wide infectivity of this emerging pathogen. Overall, all this evidence reinforces the need to establish surveillance strategies in wildlife and domestic animals to detect and characterize new adenoviruses with zoonotic potential.

### 2.5. Novel members from the phylum *Cressdnaviricota*

Within this phylum, there are seven families of Rep-encoding viruses with single-stranded, circular DNA genomes [62]. Complete genomes of members of two of these families (i.e. *Circoviridae* and *Genomoviridae* families) were identified. Circovirids, which include *Circovirus* and *Cyclovirus* genera, are associated with animal diseases, particularly the former [63,64]. In contrast, much less is known about genomoviruses, whose presence could be due to food-borne transmission [65], although their possible role in animal pathogenesis cannot be ruled out.

A genome associated with *Circoviridae* family, MAVG5, was assembled de novo in a pool of faecal samples from species *Crocidura russula*, which belongs to *Soricidae* family (pool P40) (**Figure 2** and **Tables S1** and **S4**). In this family, viruses classified into different species share less than 80% genome-wide pairwise sequence identity [66]. A BLASTn analysis of MAVG5 showed a peak sequence identity of 97.9% and a 99.0% query coverage to human-associated cyclovirus 6 strain RI46/ITA (Acc. MZ201304.1), and a ML tree of the Rep protein confirmed its classification as a cyclovirus (**Figure 6** and **Table S4**). In addition, both sequences shared 89.2% genome-wide pairwise identity, thus they could be considered variants of the same species. Although cycloviruses are thought to be associated with arthropods or contamination rather than animal infection, a potential role has been suggested in human respiratory infections [67], human enteric infections via foodborne or fecal routes of transmission, as well as a high prevalence of infection in pigs and poultry [68]. Thus, obtaining additional information on the location of cycloviruses and possible animal reservoirs may help in the future to understand their true role in the animals where they are found.

**Figure 6.**
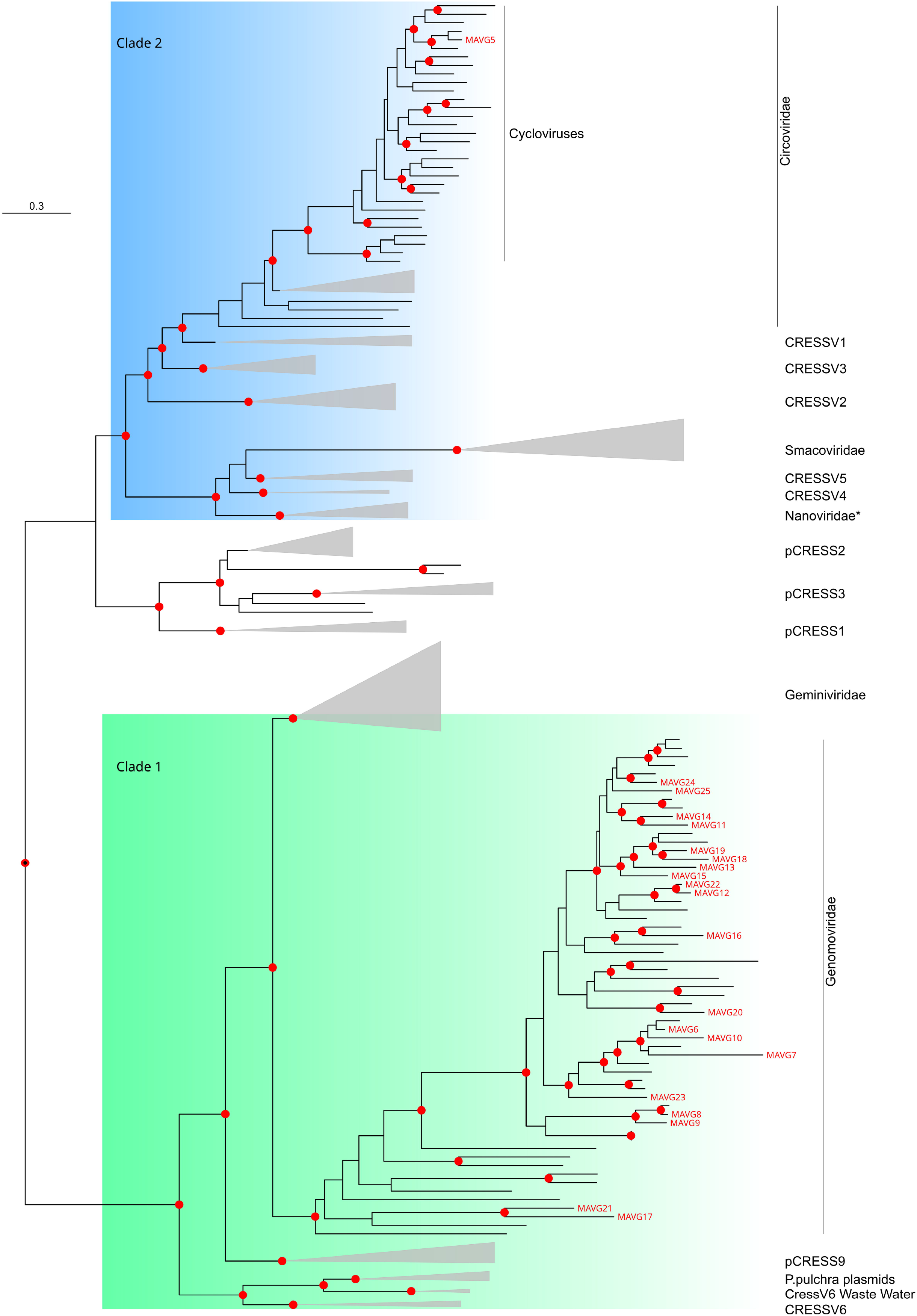
Subtree obtained from a ML tree of the Rep protein from CRESS-DNA viruses [72]. Nodes are collapsed by family or other undefined taxonomic groups, with the exception of the families *Circoviridae* and *Genomoviridae*. Novel MAVGs are labelled in red. Phylogenetic analysis was done using substitution model LG+G4. SH-aLRT and ultrafast bootstrap values higher than 80 and 95 respectively, are indicated in red circles. The scale bar indicates the evolutionary distance in amino acid substitutions per site.

Twenty MAVGs belonging to *Genomoviridae* family were detected in two fecal samples belonging to *Apodemus sylvaticus* and *Sorex coronatus*, from *Rodentia* and *Soricomorpha* orders, respectively (**Figure 2** and **Tables S1** and **S4**). Genomes sizes ranged between 2.1 and 2.4 kb, as expected for genomoviruses [65]. MAVGs showed one or two ORFs encoding the Rep protein and one ORF for the capsid protein. The genus demarcation criterion for genomoviruses is based on Rep amino acid sequence phylogeny. Accordingly, eighteen MAVGs were assigned to the *Gemycircularvirus* genus, while MAVG17 and MAVG21 were close to *Gemytondvirus* genus, although they could also be considered members of a new genus (**Figure 6** and **Figure S1**). For genomoviruses, the species delimitation threshold is a genome-wide pairwise sequence identity of 78% [65]. Based on this threshold, 16 new species were identified, one of which included two MAVGs (i.e. MAVG12 and MAVG22), whereas the remaining MAVGs (i.e. MAVG6, MAVG7, and MAVG9) could be considered strains of already reported species.

Since the initial identification of a genomovirus in a fungus [69], hundreds of genomes have been described in metagenomic studies, leading to the current definition of a total of ten genera [65]. For instance, members of this family has been reported in soil metagenomes [65], human cerebrospinal fluid [70] or bat liver samples [71]. In this scenario, characterizing and understanding the diversity of this family may shed some light on many open questions, such as its host range, ecological role or involvement in pathogenesis.

## 3. Conclusions

The zoonotic potential of DNA viruses is less than that of RNA viruses. However, there are numerous examples that this threat is real. Similarly, DNA viruses are not considered to be a likely source of future pandemics, due to the higher fidelity of their polymerases. However, there are other variability-generating mechanisms, such as recombination, which play a major role in the generation of new adenoviruses, for example by increasing their pandemic potential [73]. Metagenomic studies can help us to identify DNA viruses with zoonotic potential, although it is important to bear in mind that the ability to detect them may depend on the nature of the samples to be analysed. In the case of faecal samples, for example, the host and the collection procedure may determine the presence of inhibitory agents for virus detection. Nevertheless, metagenomics is becoming an essential tool for a better understanding of the virosphere and the establishment of surveillance strategies against potential emerging viruses.

## 4. Material and methods

### 4.1. Study area and sample collection

Box-like traps (i.e. Sherman, Ugglan, and Mesh traps) were used to capture small mammals from different habitats in eight Spanish provinces (Alicante, Cantabria, Huelva, León, Madrid, Salamanca, Valencia, and Zamora; **Figure 1**). Captures took place from March to November 2022. In total, 160 individuals were captured and the species, sex and age were identified before release, except in a few cases, where samples were taken, without capture, from burrows of previously identified species. The taxonomic orders of the captured animals included *Rodentia, Lagomorpha*, and *Eulypotyphla*. Fresh faecal samples were collected from catchment boxes, or occasionally from burrows, and kept individually in tubes containing 500 μL of 1X phosphate-buffered saline (PBS) at −20 °C until they were transported to the laboratory and stored at −80 °C for further processing.

### 4.2. Identification of samples with potential inhibitors

Preliminary sample processing yielded negative results for virus detection by massive sequencing (not shown). This suggested that some samples might contain substances involved in the degradation of nucleic acids/viral particles, or in inhibiting extraction/sequencing stages, such as urine [20] or dietary components [21]. To identify these samples with inhibitor potential, they were processed individually as previously described [23]. Briefly, an aliquot of each sample was homogenized in a Precellys Evolution tissue homogenizer (Bertin) and the supernatant obtained after centrifugation was filtered using Minisart cellulose acetate syringe filters with a 1.2 µm pore size (Sartorius). Then, each filtrate was incubated for 1h with 10^7^ PFU of a recombinant vesicular stomatitis virus in which the G protein was replaced by the green fluorescent protein. Finally, half of the volume was used to infect in triplicate a monolayer of BHK cells in 96-well plates and photographs were taken 16 hours post-infection to detect fluorescent cells using an Incucyte^®^ microscope, as previously described [75]. In addition, for some samples, nucleic acids were extracted from the other half of the filtrate volume using the QIAamp Viral RNA mini kit (Qiagen) and VSV-specific RT-qPCR was performed with three replicates per sample [74]. A sample was considered inhibitory if a reduction of fluorescence of more than 90% and/or a decrease of more than two Ct values were observed when compared to positive controls in culture assays and qPCR, respectively (Results not shown). Consequently, all samples with inhibitory potential were eliminated from the study (**Table S1**).

### 4.3. Sample processing and nucleic acids extraction

After excluding those samples with inhibitory potential, the remaining 76 samples were combined as previously described [23] into a total of 13 pools, each containing between one and 12 samples from the same species (**Table S1**). Processing of each pool was carried out as described above for individual samples and nucleic acids were obtained using the QIAamp Viral RNA mini kit (Qiagen), which is routinely used for RNA and DNA co-purification [76,77].

### 4.4. Sequencing and viral sequence detection

The preparation of libraries from extracted nucleic acids was carried out using the Nextera XT kit (Illumina) with 15 amplification cycles and paired-end sequencing was performed on a NextSeq 550 device with the read length of 150 bp at each end. Raw reads were processed with fastp 0.23.2 [78], including deduplication, quality filtering, and trimming with a threshold of 20, and those reads below 70 nucleotides in length were removed. De novo assembly was conducted using SPAdes v3.15.4 [79] and MEGAHIT v1.2.9 [80], followed by clustering with CD-HIT v4.8.1 [81] to remove redundancies. Contigs shorter than 1,000 nucleotides were discarded. Taxonomic classification was performed with Kaiju v1.9.0 [82] using the prebuilt nr database downloaded on 6 June 2023, which contains a subset of the NCBI BLAST nr database of archaea, bacteria and viruses. Viral contigs were identified using Virsorter2 v2.2.4 [83] and further analyzed with CheckV v1.0.1 [24] for genome quality assessment. Sequences associated with phages or unclassified viral families were excluded. The remaining contigs were selected based on size, completeness, and potential vertebrate infectivity. Coverage statistics for the viral contigs were determined by remapping the filtered and trimmed reads to their corresponding contigs using Bowtie2 v2.2.5 [84]. In addition, viral contigs were compared against NCBI databases using BLAST [85] to obtain sequence identity values and refine annotations. The raw sequence reads were deposited in the Sequence Read Archive of GenBank under accession numbers SRR30838292-304. The MAVGs described in this study, which corresponded to complete or nearly complete genomes, were deposited in GenBank under accession numbers PQ576916-40 (**Table S4)**.

### 4.5. Phylogenetic analysis

The search for sequences similar to each contig of interest was performed using DIAMOND v2.0.15.153 [86] with the blastp option and the NCBI nr database downloaded on April 15, 2024. For each contig, the 100 closest matches obtained from DIAMOND were analyzed to identify potential associations with viruses infecting vertebrates. Additionally, protein domains were annotated using Interproscan v5.63-95.0 [87] with the Pfam database v35.0. Open reading frames (ORFs) were predicted using ORFfinder (https://www.ncbi.nlm.nih.gov/orffinder). For contigs assigned to viruses with potential to infect vertebrates, a multiple sequence alignment was obtained from amino acid and nucleotide sequences specific to each viral family using Clustal Omega v1.2.3 [48] or MAFFT v7.505 [88], respectively. Phylogenetic analyses were performed with IQ-TREE v2.3.6 [89] and model selection was done using the built-in ModelFinder function [90]. Branch support was assessed using 1,000 ultra-fast bootstrap replicates [91] and 1,000 bootstrap replicates for the SH-like approximate likelihood ratio test.

## Supplementary materials

Figure S1: ML tree of the family *Genomoviridae* based on 94 representative amino acid sequences of Rep gene plus 24 genomovirus Rep sequences recently reported in a metagenomic study of bat faeces in Spain [23]. Taxa are denoted by GenBank accession number and virus name, and novel MAVGs are labelled in red, indicating with an asterisk those that are defined as new species. When a new species includes more than one novel MAVG, only one is indicated with an asterisk. Phylogenetic analysis was done using substitution model Q.pfam+F+I+G4. SH-aLRT and bootstrap values higher than 80 and 95, respectively, are indicated with red circles. The tree is rooted at midpoint and the scale bar indicates the evolutionary distance in amino acid substitutions per site.

Table S1: Small mammal species, collection sites, pool distribution, and inhibitory potential of each sample.

Table S2: Illumina reads and number of viral contigs obtained.

Table S3: List of viral contigs classified by CheckV as high quality or complete genome.

Table S4: Descriptions, accession numbers, and proposed names for the novel MAVGs.

Table S5: Genome information for papillomavirus MAVGs.

Table S6: Genome information for MAVG3 polyomavirus.

## Funding

This research was funded by grant PID2020-118602RB-I00 from the Spanish Ministerio de Ciencia e Innovación (MICINN) and cofinanced by FEDER funds, and grant

CIAICO/2022/110 from the Conselleria de Educación, Universidades y Empleo (Generalitat Valenciana).

## Institutional Review Board Statement

Samples consisted of faeces from wild animals captured using box-like traps or collected without handling or trapping specimens. After visual identification of the individuals, they were released and a faecal sample was collected from the trap. According to the European directive regulating the protection of animals used for scientific purposes (2010/62/EU, Article 1), subsequently transposed into Spanish legislation (Royal decree 53/213, 1 February, Article 2), procedures used in this study (i.e. capture, non-invasive handling and in situ release of wild animals) are not subject to the condition of animal experimentation and, therefore, an IACUC approval document is not required, but specifically a permit from the competent regional.

## Acknowledgments

We would like to thank the collaboration of different research groups and organizations that have supported the fieldwork, in special to Dr. Francisco Carro (EBD), Dr. Marius V. Fuentes (UV), Dr. Sandra Saiz-Elipe (Rifcon), Ayuntamiento de Madrid, and Fundación Naturaleza y Hombre (Cantabria).

