## Supplementary figures and images for "Complete genomes of DNA viruses in faecal samples from small terrestrial mammals in Spain"

### Figure S1

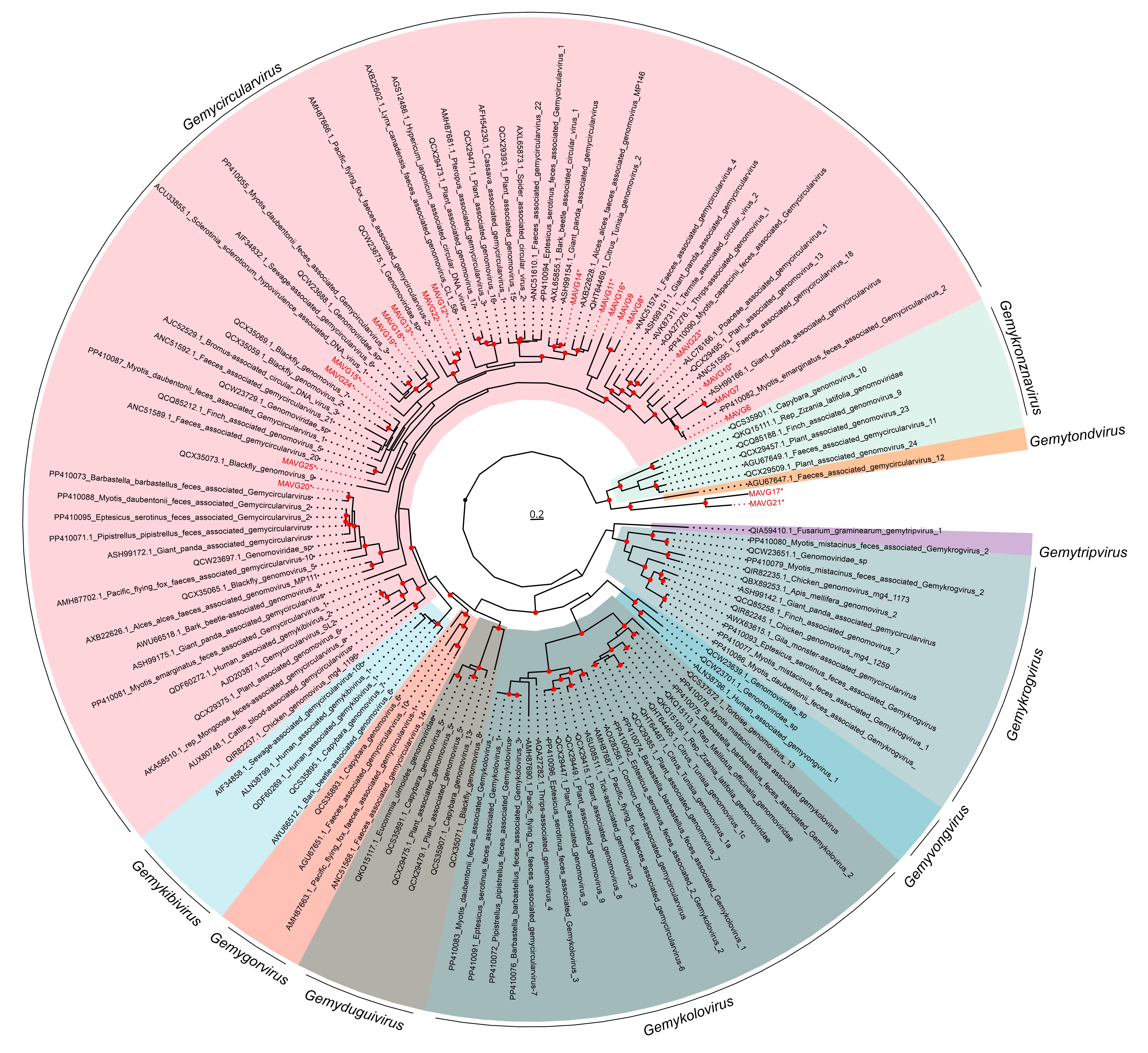
